# Synthetic neuromelanin as a trigger of inflammation in the brain – new mouse model of Parkinson’s disease

**DOI:** 10.1101/2023.04.28.538536

**Authors:** Anna Tejchman-Skrzyszewska, Martyna Strzelec, Marta Kot, Paula Chlebanowska, Bogna Badyra, Małgorzata Sobocińska, Luigi Zecca, Marcin Majka

**Affiliations:** Department of Transplantation, Institute of Pediatrics, Jagiellonian University Medical College, Wielicka 265, Cracow, 30-663, Poland; Institute of Biomedical Technologies – National Research Council of Italy, 20090 Segrate (Milano), Italy

## Abstract

Parkinson’s disease (PD) is a neurodegenerative disease that is an increasing threat to an aging society. The idiopathic form of PD accounts for over 90% of all cases, and the current etiology is still unknown. One of the reasons hindering research on this form of PD is the lack of an appropriate animal models. Among mouse models of the disease, those based on the administration of neurotoxins such as 1-methyl-4-phenyl-1,2,3,6- tetrahydropyridine (MPTP) or 6-hydroxydopamine (6-OHDA) to the substantia nigra pars compacta (SNpc) or striatum are predominantly used. In these models, there are metabolic disturbances causing oxidative stress in the SNpc or striatum, which ultimately leads to the death of dopaminergic neurons. However, the models used so far have serious limitations, most of all they do not fully reflect the processes occurring in the course of the disease and do not consider the involvement of inflammation in the etiology and pathogenesis of PD. In this study we show that the administration of synthetic neuromelanin, which activates astrocytes and microglia, induces the inflammation and may be involved in degeneration of dopaminergic neurons. Neuromelanin under physiological conditions acts as a neuroprotector, however, released from dying dopaminergic neurons is an important factor activating astrocytes, microglia and causing neuroinflammation. Since one of the causes of Parkinson’s appear to be the death of dopaminergic neurons overloaded with neuromelanin and consequent pathological activation of microglia, the use of synthetic neuromelanin reflect the natural pathological processes occurring during the development of the disease.

## Introduction

Parkinson’s disease (PD) is serious clinical problem affecting millions of people worldwide (1,2). Despite a lot of efforts, the effective treatment has not been developed yet (2). One of the reasons could be lack of animal models that resemble in details the course of the disease (3). Among the most commonly used models are mouse models of PD that based on the administration of neurotoxins such as 1-methyl-4-phenyl-1,2,3,6-tetrahydropyridine (MPTP) or 6-hydroxydopamine (6-OHDA) to the striatum or rotenone in the gastric form (4). These compounds cause acute metabolic toxicity by inducing the formation of oxidative stress in the *substantia nigra* of midbrain, which in effect leads to the death of dopaminergic neurons.

However, these models have serious limitations, above all they do not fully reflect the processes occurring in the course of the disease (5,6). Under physiological conditions, the process protecting dopaminergic neurons from damage caused by oxidation of catecholamine neurotransmitters, particularly dopamine, which leads to the accumulation of reactive quinone in the cytosol occurs (7). This process is the formation of neuromelanin, a dark pigment that accumulates during aging (8,9,10).

Neuromelanin accumulates in the cytoplasmic organelles surrounded by a double membrane in various areas of the brain but mainly in dopaminergic neurons in the *substantia nigra pars compacta (SNpc)* (10, 11). Neuromelanin also protects mDA by binding heavy metals with particular affinity to iron ions (11, 12, 13). Although neuromelanin acts as a neuroprotector under physiological conditions, released from dying dopaminergic neurons is a key factor in activating microglia (14, 15, 16).

Microglia are the innate immune cells that maintain the CNS homeostasis under physiological conditions but over-activated may produce pro-inflammatory cytokines such as interleukin (IL)-1β, interleukin (IL)-6 and tumor necrosis factor (TNF) 5,6 (nature scientific reports) (16, 17). Microglia, the first line of defense reacts external or internal signals by producing large amounts of cytokines e.g., IL-1β which in turn convert astrocytes into a neurotoxic A1 phenotype in a range of neurological diseases (18, 19), activating them to produce neurotoxic factors (17). Taking together, microglia with astrocytes orchestrate production of proinflammatory cytokine cocktail causing the death of tyrosine hydrolase positive dopaminergic neurons (20). Known mediators promoting progression of PD such as LPS induce the transcription factors as MyD88 or Trif via toll-like receptor (TLR)4 and activate downstream signals such as NF-κB (21, 22).

Our goal was to create a new mouse model of PD based on the activation of microglia with synthetic neuromelanin and the development of neuroinflammation. The study assumes the use of two newly synthesized neuromelanin derivatives (Pheo-β-LG and Pheo-β-fLG) based on the conjugation of DA-melanin with a fibrillar protein, β-lactoglobulin (obtained as part of international cooperation from Prof. Luigi Zecca from the Institute of Advanced Biomedical Technologies -CNR, Segrate, Milan, Italy), mimicking the structure of neuromelanin naturally occurring in the brain (7). Because one of the causes of PD appears to be the death of dopaminergic neurons overloaded with neuromelanin and, as a consequence, pathological activation of microglia, the use of synthetic neuromelanin reflects processes occurring during the development of the disease (7, 23, 24, 25). In present study hypothesis about the pro-inflammatory role of synthetic neuromelanin in the course of PD was investigated (7). Importantly, new mouse model of PD reflecting well some essential aspects of PD development and progression may be used to better understand the diseases and to test the possibility of rebuilding the neural network by administering cell therapeutics based e.g., on dopaminergic neurons obtained by differentiating induced pluripotent stem cells.

## Materials and methods

### Mice

Male 8–12-week-old NOD-SCID mice (NOD.Cg-Prkdcscid/J, Charles River Laboratories) were used in all experiments. Mice were maintained under specific pathogen-free conditions and a standard 12 h light/ dark cycle with unrestricted access to food and water. Mice were housed in individually ventilated cages in groups of up to five mice and were acclimatized for a minimum of 1 week prior to procedures. All procedures involving live animals were approved by The Animal Welfare and Ethics Committee.

### Stereotactic injection of neuromelanin, 6-OHDA and PBS

Injections of a mixture of the two types of synthetic neuromelanin (NM) into SNpc at a concentration of (0.85 mg/ml of Pheo-β-LG and 0.85 mg/ml of Pheo-β-fl) 3μl were done during the stereotaxic surgery. Adult male mice of the NOD-SCID strain (∼30–35 g) were administered 2 µl of 6-hydroxydopamine (6-OHDA, Sigma) dissolved in PBS at a concentration of 3.75 µg/µl with the addition of 0.01% ascorbic acid at the rate of 0.5 µl/min during a stereotaxic surgery. PBS was injected in an amount of 3μl at the rate of 0.5 µl/min during a stereotaxic surgery. Mice were anesthetized by intraperitoneal administration of ketamine (87.5 mg/kg body weight) and xylazine (12.5 mg/kg).

### Measurement of spontaneous activity in cylinder test

The animals were individually placed in a 13 cm diameter glass cylinder and spontaneous activity was recorded for 3 min (29, 30, 31). The cylinder was situated on a piece of glass with a mirror positioned at an angle beneath the cylinder to allow the camera for a clear view of the animal movements along the ground as well as along the walls of the cylinder. The animals were allowed to move freely while being recorded with the camera and quantification of forelimb use was done by an observer blinded to the animal group identity (32). Forelimb use was defined as a vertical movement (rearing) with the simultaneous placement of both palms on the cylinder wall. Touches by a single palm were not calculated. Number of forelimbs uses was compared for wild – type, PBS, 6-OHDA and neuromelanin mice at 1, 2, 3 and 4 weeks after surgery and for PBS and neuromelanin group at week 1 and week 27 after the surgery.

### Extraction of RNA and Reverse Transcription Reaction

Isolation of mRNA was made by GeneMATRIX Universal RNA/miRNA Purification Kit (EURx), according to the manufacturer’s instruction. RT-PCR with random primers (Promega, Madison, WI, USA) and Moloney murine leukemia virus M-MLV reverse transcriptase (Promega, Madison, WI, USA) was performed according to the vendor’s protocol. Frozen in liquid nitrogen immediately after sacrificing the mice and after dissecting the tissue, the SNpc brain sections was treated using the QIAzol Lysis Reagent (Qiagen) for lysis of fatty tissues before RNA isolation. Sections were then homogenized with a TissueLyser II, an automatic homogenizer (Qiagen) according to the manufacturer’s instructions. Purification of total RNA from fatty tissues as brain was performed using the RNeasy Mini Kit (Qiagen) according to the manufacturer’s instructions. RT-PCR with random primers (Promega, Madison, WI, USA) and Moloney murine leukemia virus MMLV reverse transcriptase (Promega) was performed according to the vendor’s protocol.

### Real-Time PCR

Gene expression levels were evaluated using quantitative real-time PCR analysis (qPCR) and Quant Studio 7 Flex System (Applied Biosystems, Foster City, CA, USA). The following TaqMAN probes were used: TH (Hs00165941_m1), GAPDH (Hs02758991_g1), Il1b (Mm00434228_m1), Il6 (Mm00446190_m1), Tnf (Hs01547104_g1), Myd88 (Mm00440338_m1), Nfkb1 (Mm00476361_m1), Trem1 (Mm01278455_m1), Trem2 (Mm04209424_g1), Iba1 (Mm00494477_m1), GAPDH (Mm99999905_m1), TH (Mm00447557_m1) and Blank qPCR Master Mix (2×) (EURx). The mRNA expression levels for all samples were normalized to the levels of housekeeping gene GAPDH using the 2−ΔCt method, which allowed us to calculate the relative expression of genes.

### BV-2 Microglia Culture

A mice BV-2 cell line was obtained from ATCC (USA). BV-2 cells were cultured in DMEM (Invitrogen) supplemented with 5% v/v fetal bovine serum (Eurx, Gdansk, Poland) and 1% penicillin/streptomycin (Thermo Fisher Scientific) at 37°C in a humidified incubator under 5% CO2 and 95% humidity. Cells were seeded at a density of 2.5 × 10^5^ cells/ml 24 h before stimulations at which point cells were >90% confluent. Prior to stimulation, cells were washed twice with DPBS (Invitrogen) and cultured in serum-free DMEM for at least 1h. Cells were stimulated for 4 h with one of the following: LPS (E. coli 0127:B8 100 ng/ml; Sigma-Aldrich); solutions of mix of synthetic neuromelanins (NM 10 μg/mL of Pheo-β-LG and 10 μg/mL of Pheo-β-fl). Cells were passaged with Accutase cell detachment solution (BioLegend, San Diego, CA, USA) and seeded on new dishes. Cells were routinely tested for mycoplasma contamination using a MycoAlert™ Mycoplasma Detection Kit (Lonza).

### Immunofluorescent Staining

The brains were isolated, washed in DPBS and then fixed in 4% paraformaldehyde for 24 h at room temperature. After incubation, they were stored up to one week in DPBS at 4°C or embedded into paraffin blocks and then 3 µm slides were prepared in the Pathology Laboratory of the Children Hospital in Cracow. From brains stored in DPBS 50 µm slides of the coronal brain sections were prepared (Vibratome). The 3 µm slides were rinsed with TBS containing 0.025% Triton X-100 (Sigma–Aldrich) twice for 5 min and they were then were blocked in 1% bovine serum albumin (BSA, Sigma–Aldrich) in tris-buffered saline (TBS, 50 mM Tris-Cl, pH 7.6; 150 mM NaCl) for 1 h at room temperature. Subsequently, the slides were incubated with appropriate primary antibody diluted in 1% BSA in TBS for 1 h at 4°C. Antibodies were as follows: mouse anti-tubulin antibody, beta III isoform (Tuj1 MAB1637, Sigma–Aldrich), rabbit anti-tyrosine hydroxylase antibody (TH AB152, Merck-Millipore, CA, USA). The slides were washed with TBS containing 0.025% Triton X-100 twice for 5 min and were incubated with Hoechst (Sigma–Aldrich) and secondary goat anti-rabbit or anti-mouse antibodies that were conjugated with Alexa Fluor 555 (Thermo Fisher Scientific) or Alexa Fluor 488 (Thermo Fisher Scientific) diluted in 1% BSA in TBS for 1 h at room temperature in the dark.

### TEM imaging

The brain tissue was fixed in 2.5% glutaraldehyde at room temperature for 24h. The cells were fixed in a monolayer with 2.5% solution of glutaraldehyde in PBS with Ca^2+^ and Mg^2+^ (pH 7.3) for 3h at room temperature, then scraped with a cell scraper. Next the tissue and cell pellets were covered with fresh glutaraldehyde solution and stained in 1% solution of osmium tetroxide. In the next steps, material was rinsed and dehydrated in ethanol series (50%, 70%, 90%, 95%, 100%) and methanol, and embedded in epoxy resin Epon 812 (Serva, Germany). The ultrathin sections (90 nm thick) were contrasted with 2% uranyl acetate and lead citrate and analyzed and photographed under a Jeol JEM 2100 transmission electron microscope at 80 kV.

### Induced Pluripotent Stem cells differentiation

Induced Pluripotent Stem cells (iPSCs) were generated from Peripheral Blood Mononuclear cells (PBMCs) isolated from healthy volunteer and PD patients. Isolation of PBMC, reprogramming and cell culture were described by Chlebanowska *et al.* (3). The study was approved by Jagiellonian University Bioethical Committee in Kraków, decision number KBET/173/B/2012. Dopaminergic neurons differentiation was performed as describe by Krisk et al. (33). BioLegend, San Diego, CA, USA) and seeded at density 3,5 x 105 cells/cm2. From day 20 medium was consisted of Neurobasal Plus, B27 Plus supplement, BDNF (brain-derived neurotrophic factor, 20 ng/ml; Peprotech, London, United Kingdom), ascorbic acid (AA; 0.2 mM, Sigma-Aldrich, St. Louis, MO, USA), GDNF (glial cell line-derived neurotrophic factor, 20 ng/ml; Peprotech), TGFβ3 (transforming growth factor type β3, 1 ng/ml; Peprotech), dibutyryl cAMP (0.5 mM; Sigma-Aldrich), 100 U/ml Penicillin/Streptomycin (Thermo Fisher Scientific) and DAPT (10 nM; Stemgent, Cambridge, MA). Cells were stimulated with either LPS (100 ng/ml; Sigma-Aldrich) or with NM (10 μg/mL of Pheo-β-fLG and 10 μg/mL; kindly provided by Dr. Luigi Zecca Institute of Biomedical Technologies – National Research Council of Italy, Milano, Italy) by two weeks, one week, 3 days and 24 hours till day 35. The LPS and NM were added to fresh medium. Fresh medium was changed every day. The control cells were collected on day 35. Cells were cultured in a humidified atmosphere of 5% CO2 in 37°C. After stimulation, cells were washed by PBS and collected for future analysis.

## Results

### Neuromelanin similarly to LPS activate BV-2 microglial cells to produce proinflammatory cytokines (IL-1, IL-6 and TNF-***α***) in NF-***κ***B and MyD88 dependent manner

One of the goals of the project was to check whether synthetic neuromelanin induces inflammation in the microglial cells, which in turn leads to damage to dopaminergic neurons in the *SNpc* in the brain of mice.

Ultrastructural (TEM) analysis of BV-2 microglia cells treated with synthetic neuromelanins solution revealed numerous granules of electron dense material identified as neuromelanin (Fig. 1a). Black pigment granules internalized by BV-2 have been localized in single membrane-bound compartments (Fig. 1a). In cytoplasm of control cells (standard cultured cells) similar structures were not observed. Therefore, as a preliminary experiment, an *in vitro* check was performed whether synthetic neuromelanin induces the expression of pro-inflammatory cytokines in the BV-2 murine microglia cell line to a degree similar to that of the LPS molecule. An analysis of gene expression levels reflected significant upregulation of pro-inflammatory cytokines shown as a relative change in real-time PCR (qPCR) analyses in BV-2 microglia cells treated with solutions of mix of synthetic neuromelanins (NMs) 10 μg/mL of Pheo-β-LG and 10 μg/mL of Pheo-β-fl or LPS 100 ng/ml for 4 h visualized in Figure 1b. Relative changes in real-time qPCR analyses in microglia cells are reported for IL-1β, IL-6 and TNF-α (Fig. 1b). In addition to pro-inflammatory cytokines, gene expression analysis was performed for the triggering receptor expressed on myeloid cells (TREM) family which are cell surface innate immune receptors of the immunoglobulin superfamily expressed on myeloid cell populations throughout the body, including microglia in the brain (22, 26, 27). Previous studies show that microglia expression of Trem1 and Trem2 genes is counter-regulated by lipopolysaccharide (LPS) *in vitro* in both adult mouse and human microglia (22). We obtained similar results for LPS and neuromelanin both causing significant TREM-1 induction and TREM-2 suppression in the BV-2 cell line (Fig. 1b). Our data show also that both neuromelanin and LPS significantly induce NF-κB expression. Neuromelanin induced expression a **∼**4 fold and LPS a **∼**6 fold, with LPS inducing NF-kB expression a **∼**1,5 fold more than neuromelanin (Fig. 1b). This indicates NF-κB as a common signaling intermediary controlling divergent in vitro responses for TREM-1 and TREM-2. Our data shows coordinated but divergent regulation of TREM microglial receptor expression with a central role for NF-κB. Neuroinflammatory states that alter the balance of TREM expression may therefore have a significant effect on microglial inflammatory activity with implications for neuroinflammatory and neurodegenerative diseases. LPS activates signaling dependent on Toll-like receptor 4 (TLR4), myeloid differentiation factor 88 (MyD88) which is evidenced by a significant increase in MyD88 ∼2.125-fold expression after LPS stimulation in the BV-2 cell line (Fig. 1b). Neuromelanin increases the expression of MyD88 but this increase is not significant *in vitro*. Our results suggest that TREM-1 activation in BV-2 cell line may be associated with TLR4-MyD88-NF-κB-dependent signaling.

**Figure 1.**
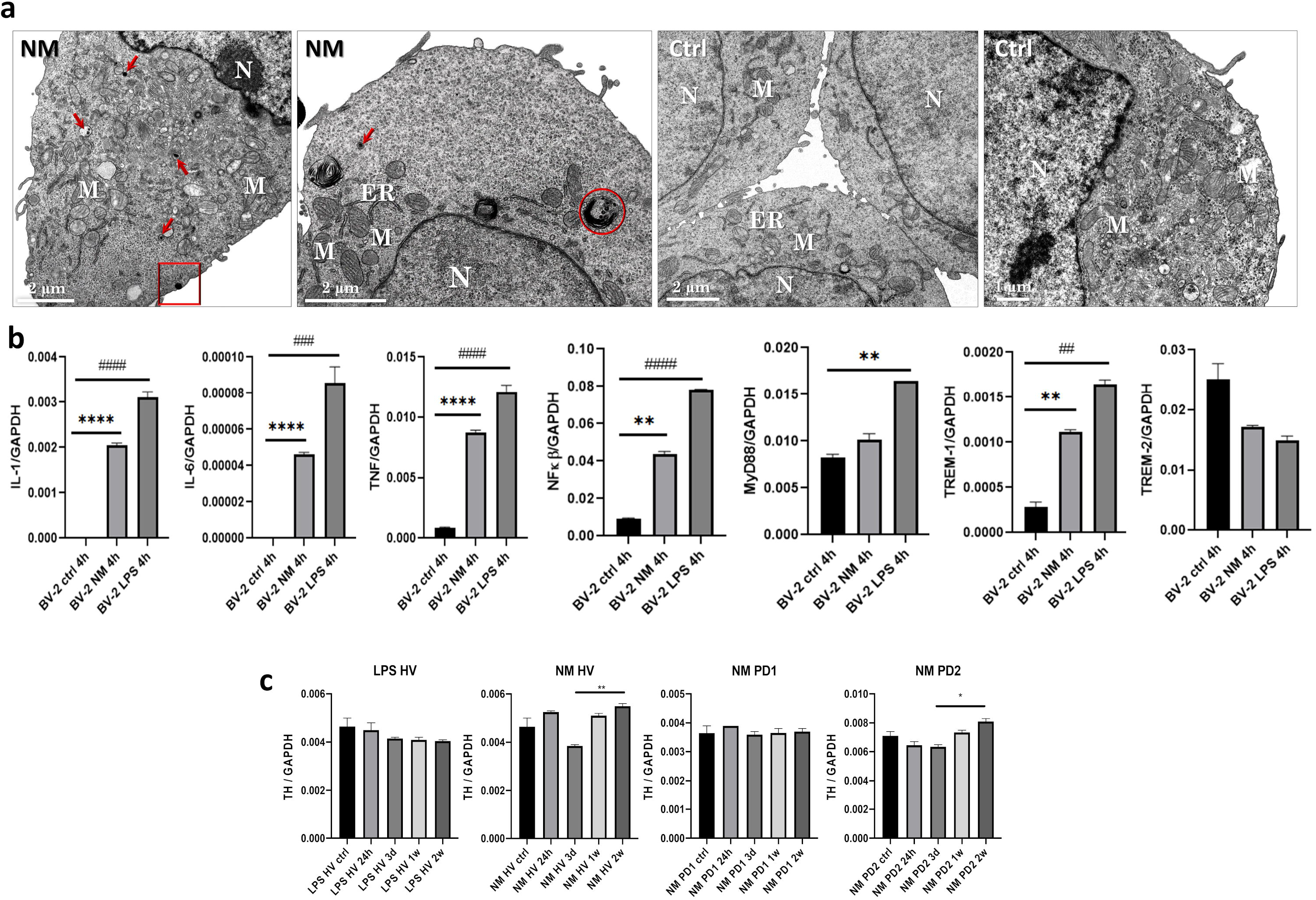
Activation of BV-2 and upregulation of tyrosine hydroxylase in iPS derived dopaminergic neurons by synthetic neuromelanin. **a)** Ultrastructure of murine microglial cells (BV-2). Neuromelanin uptake in BV-2 cells (NM); Ctrl – control cells, N – nucleus, M – mitochondria, ER – endoplasmic reticulum, red arrows, red circle – neuromelanin granules; Red square – internalization of neuromelanin into BV-2 cell; TEM, bar=2µm or 1µm respectively. **B)** Relative changes in qPCR analyses in BV-2 microglia cells treated with solutions of mix of synthetic neuromelanins (NM 10 μg/mL of Pheo-β-LG and 10 μg/mL of Pheo-β-fl) or LPS 100 ng/ml for 4 h. Relative changes in qPCR analyses in microglia cells are reported for IL-1β, IL-6, TNF-α, NFκB, MyD88, TREM-1 and TREM-2. LPS – lipopolysaccharyde; GAPDH - glyceraldehyde-3-phosphate dehydrogenase; IL-1β – interleukin 1β; IL-6 – interleukin 6; TNF- α – tumor necrosis factor; NF-κB – nuclear factor kappa-light-chain-enhancer of activated B cells; MyD88 – Myeloid differentiation primary response 88; TREM-1 – Triggering receptor expressed on myeloid cells 1; TREM-2 – Triggering receptor expressed on myeloid cells 2. Relative expression of all genes to GAPDH were validated by qPCR. Student t-test was performed to verify statistical relevant between the compared groups in selected time point. The graphs show results from two different wells. The data represent the mean ± SEM. ** p < 0.01; *** p < 0.001; **** p < 0.0001. **c)** Expression (RT-qPCR) of tyrosine hydroxylase (TH) in dopaminergic neurons derived from healthy volunteer and PD patients treated with neuromelanin (NM) or lipopolysaccharides (LPS). During differentiation the generated neurons were treated with NM or LPS for 24h, 3days (3d), 1 week (1w), 2 weeks (2w). In neurons from healthy volunteer treated with LPS, level of TH expression does not significantly change. In neurons from healthy volunteers treated NM, TH expression shows significant difference between groups on day 3 and 2 weeks. ** *p* < 0.005. The expression of TH is on the same level during NM exposition in PD1 neurons. The expression of TH is significantly higher in 2 weeks versus 3 days in neurons from PD2 treated NM. * *p* < 0.05. TH - tyrosine hydroxylase; GAPDH - glyceraldehyde-3-phosphate dehydrogenase. Relative expression of all genes to GAPDH were validated by RT-qPCR. Student t-test was performed to verify statistical relevant between the compared groups in selected time point. The graphs show results from two different wells. The data represent the mean ± SEM.

### Synthetic neuromelanin, in contrast to LPS, changes the expression of tyrosine hydroxylase in dopaminergic neurons differentiated from iPS cells derived from healthy volunteers (HV) and patients suffering from Parkinson’s disease (PD)

In order to established the influence of synthetic neuromelanin on iPS-derived dopaminergic neurons we treaded these neurons with synthetic neuromelanin (NM) or lipopolysaccharides (LPS) as a control for 24 h, 3 days, 1 or 2 weeks and at day 36 the level of tyrosine hydroxylase (TH) was evaluated by RT-qPCR (Fig. 1c). LPS treatment of neurons from healthy volunteers did not significantly affect the level of TH. Interestingly, expression of TH in NM treated neurons from healthy volunteers significantly differs between groups treated for day 3, 1 week and 2 weeks. We observed downregulation of TH at day 3 and subsequent rise of TH expression at week 1 and 2 to levels similar to non-treated cells. Expression of TH in NM treated neurons differ between PD patients. In patient 1 (PD1) synthetic neuromelanin did not influence the TH expression. In contrast, in PD2 the expression level dropped after 24h and 3 days of NM treatment and subsequently rise at week 1 and 2 to levels similar to non-treated cells (Fig. 1c).

### Neuromelanin and 6-OHDA activates in-vivo expression of pro-inflammatory cytokines (IL-1***β***, IL-6 and TNF-***α***), microglial markers (Tmem119, F4/F80, Iba-1 and CD68), triggering receptor expressed on myeloid cells (TREM-2) leading to deceased expression of TH in NF-***κ***B and MyD88 dependent manner

We investigated the kinetics of inflammation induced by synthetic neuromelanin via analysis of the expression of pro-inflammatory cytokine genes: IL-1β, IL-6, TNF-α, as well as transcription factors: NFκB and MyD88. Another examined panel of gene expression were microglia markers: Iba-1, CD68, F4/F80, Tmem119 and receptor associated with potentiation of inflammation but also cell survival and phagocytosis: TREM-2. The mRNA expression of individual genes was measured at time points 24h, 1 week (1W), two weeks (2W) and four weeks (4W) from tissue extracted from mice injected with of NM, 6-OHDA or PBS. Schematic representation of injection time points is shown in Figure 2a. Changes in gene expression were noted after injection of 6-OHDA and NM and they were not observed after PBS administration (Figs. 2b, 3, 4). 24 hours after injection of 6-OHDA and NM, onset of the acute inflammation was recorded (Fig. 2b). Strong increase in IL-1β expression (p = 0.0023 for 6-OHDA; p = 0.0006 for NM) was accompanied by a lack of IL-6 expression and no increase in expression for TNF-α as well as NFκB (Fig. 2b). However, MyD88 expression was significantly increased after injection of 6-OHDA (p = 0.0007), which was not observed in case NM (Fig. 2b). This result corresponds to the observation at the mRNA level for Iba-1 24h after injection where the expression increased of this marker can be observed mainly after injection of 6-OHDA, but also NM but it was not clearly noticeable after PBS administration (Fig. 3b). 24 hours after the injection of 6-OHDA and NM, we also observed a sharp increase in the expression of microglial marker genes such as Tmem119 (p = 0.0186 for 6-OHDA; p = 0.0186 for NM), F4/F80 (p = 0.0002 for 6-OHDA; p = 0.0007 for NM), and also activated microglia: Iba-1 (p = 0.0002 for 6-OHDA; p = 0.0002 for NM), CD68 (p = 0.0159 for 6-OHDA; p = 0.0286 for NM) and TREM-2 receptor (p = 0.0148 for 6-OHDA; p = 0.0012 for NM) (Fig. 2b). At this point, we observe the beginning of inflammatory response. During the first week after injection of 6-OHDA or NM, expression of IL-6 occurs together with simultaneous increase in IL-1β expression (p = 0.0002 for 6-OHDA; p <0.0001 for NM) and increase in TNF-α expression (p = 0.0076 for 6-OHDA; p = 0.0283 for NM) (Fig. 3a). The expression of transcription factors is also increasing, MyD88 (p <0.0001 for 6-OHDA; p = 0.0091 for NM) and NFκB (p = 0.0012 for NM) and the marker of dopaminergic neurons decreases, TH (p <0.0001 for 6-OHDA and NM) (Fig. 3a) which indicates damage to these cells after administration of 6-OHDA and NM. The formation of a chronic inflammatory condition also begins to reveal at the protein level. The astrocyte marker GFAP appears in the area of the hippocampus and SNpc after administration of 6-OHDA and NM (Fig. 4a). Significant increase in the expression is also observed for the factors Tmem119 (p = 0.0064 for 6-OHDA; p <0.0001), F4/F80 (p = 0.0037 for 6-OHDA; p 0.0086 for NM) and CD68 (p = 0.0005 for 6-OHDA; p <0.0001 for NM) after administration 6-OHDA and NM (Fig. 3a). After administration of 6-OHDA, the expression of Iba-1 (p <0.0001) and TREM-2 (p = 0.0058) also increases and this effect is not observed after NM injection (Fig. 3a). 4 weeks after the injection, 6-OHDA and NM are observed still high expression of IL-1β (p <0.0001) (Fig. 3b). Expression of IL-6 (p = 0.0028), F4/F80 (p <0.0001 for 6-OHDA; p = 0.0007 for NM), CD68 and TREM-2 (p <0.0001 for 6-OHDA; p = 0.0004 for NM) increased significantly after NM injection (Fig. 3b). Interestingly, a significant decrease in TH expression was observed only after administration of 6-OHDA (p = 0.0007) (Fig. 3b) and not observed after NM administration. The same trend was observed at the protein level, the immunofluorescence staining showed a clear lack of TH expression in the side of 6-OHDA administration accompanied by acute astrocytosis visible by GFAP staining (Fig. 4a, b). Acute astrocytosis was also observed after NM administration, however, it did not correlate with significant decrease in the number of TH positive neurons (Fig. 4b).

**Figure 2.**
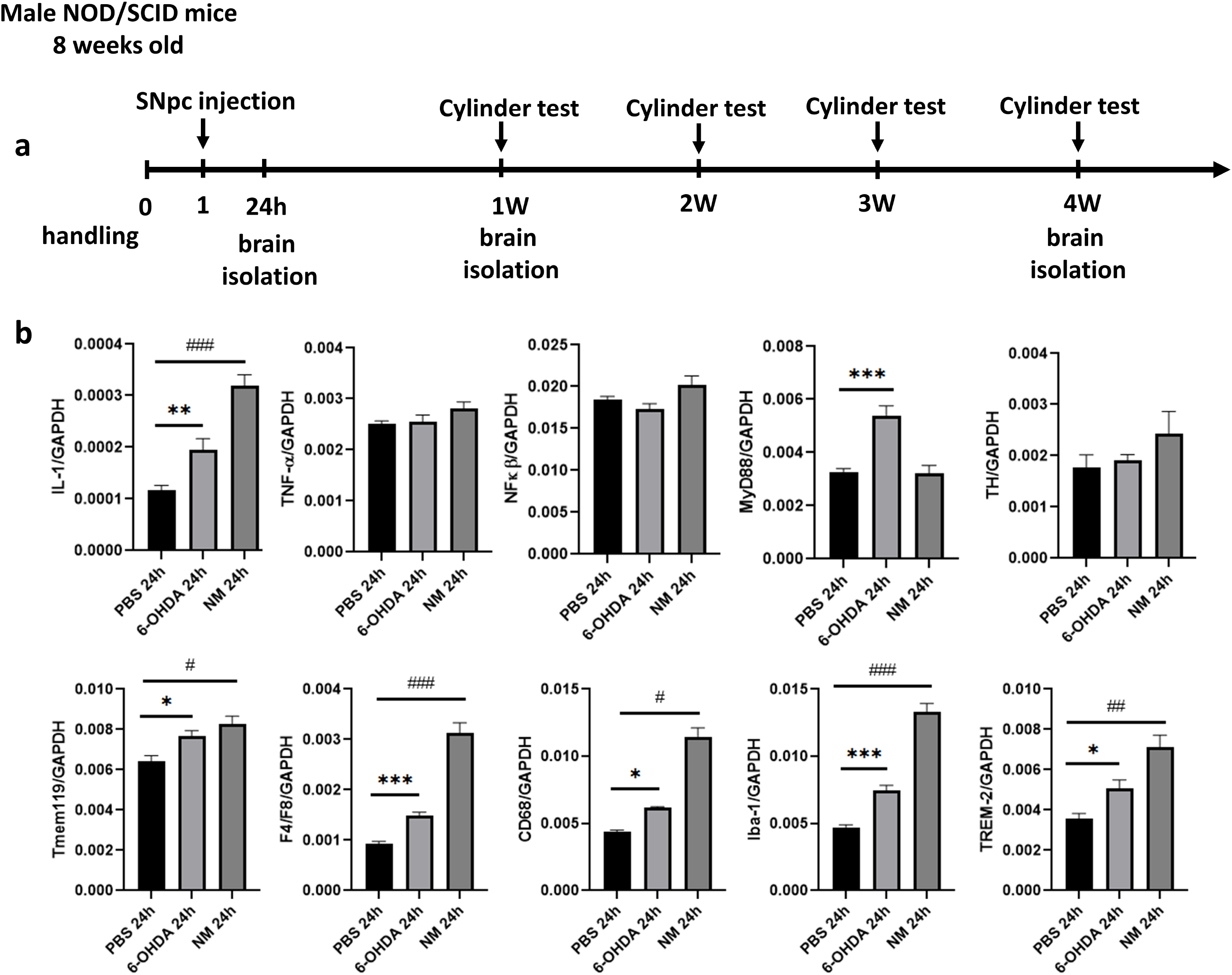
Synthetic neuromelanin activates expression of pro-inflammatory cytokines, microglial markers and TREM-2 in short term assay. **a)** Schematic representation of injection time points. **b)** Kinetics of inflammation. Relative changes in RT-qPCR analyses in brain tissue (SNpc area of left hemisphere) treated with solutions of mix of synthetic neuromelanins (3µl NM 0.85 mg/ml of Pheo-β- LG and 0.85 mg/ml of Pheo-β-fl), 6-OHDA 8 μg for 24 h. Relative changes in qRT-PCR analyses are reported for IL-1β, TNF-α, NFκB, MyD88, TH, Tmem119, F4/F80, CD68, Iba-1, TREM-2. GAPDH – glyceraldehyde-3-phosphate dehydrogenase; IL-1β – interleukin 1β; TNF-α – tumor necrosis factor; NF-κβ – nuclear factor kappa-light-chain-enhancer of activated B cells; MyD88 – Myeloid differentiation primary response 88; TH – tyrosine hydroxylase; Tmem119 – osteoblast Induction factor; CD68 – cluster of differentiation 68; Iba-1 – allograft inflammatory factor; F4/F80 – EGF-like module-containing mucin-like hormone receptor-like 1; TREM-2 – triggering receptor expressed on myeloid cells 2. Relative expression of all genes to GAPDH were validated by RT-qPCR. Mann Whitney U-test was performed to verify statistical relevant between the compared groups in selected time point. The graphs show results from two different wells. The data represent the mean ± SEM. * p < 0.05; ** p < 0.01; *** p < 0.001.

**Figure 3.**
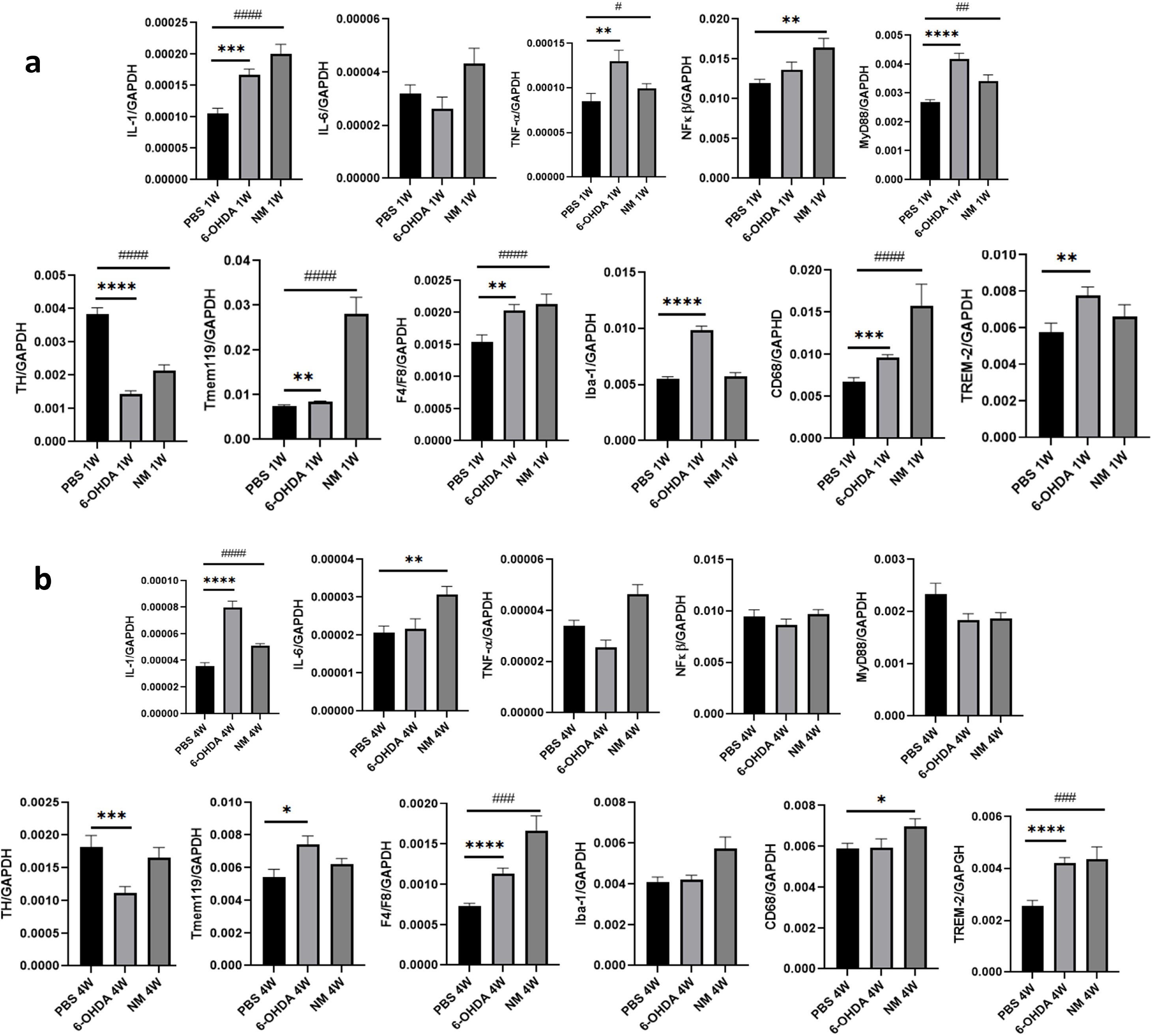
Synthetic neuromelanin activates expression of pro-inflammatory cytokines, microglial markers and TREM-2 in long term assay. **a)** Cytokines and transcription factors mRNA expression. Relative changes in RT-qPCR analyses in brain tissue (SNpc area of left hemisphere) treated with solutions of mix of synthetic neuromelanins (NM 0.85 mg/ml of Pheo-β-LG and 0.85 mg/ml of Pheo-β-fl), 6-OHDA 8 μg for 1 week (1W). Relative changes in qRT-PCR analyses are reported for IL-1β, IL-6, TNF-α, NFκB, MyD88, TH, Tmem119, F4/F80, Iba-1, CD68, TREM-2. Relative expression of all genes to GAPDH were validated by RT-qPCR. Mann Whitney U-test was performed to verify statistical relevant between the compared groups in selected time point. The graphs show results from two different wells. The data represent the mean ± SEM. * p < 0.05; ** p < 0.01; *** p < 0.001; **** p < 0.0001. **b)** Neuronal and microglial markers mRNA expression. Relative changes in RT-qPCR analyses in brain tissue (SNpc area of left hemisphere) treated with solutions of mix of synthetic neuromelanins (NM 0.85 mg/ml of Pheo-β-fLG and 0.85 mg/ml of Pheo-β-fl), 6-OHDA 8 μg for 4 weeks (4W). Relative changes in qRT-PCR analyses are reported for IL-1β, IL-6, TNF-α, NFκB, MyD88, TH, Tmem119, F4/F80, Iba-1, CD68, TREM-2. Relative expression of all genes to GAPDH were validated by RT-qPCR. Mann Whitney U-test was performed to verify statistical relevant between the compared groups in selected time point. The graphs show results from two different wells. The data represent the mean ± SEM. * p < 0.05; ** p < 0.01; *** p < 0.001. GAPDH – glyceraldehyde-3-phosphate dehydrogenase; IL-1β – interleukin 1β; IL-6 – interleukin 6; TNF-α – tumor necrosis factor; NF-κB – nuclear factor kappa-light-chain-enhancer of activated B cells; MyD88 – Myeloid differentiation primary response 88; TH – tyrosine hydroxylase; Tmem119 – osteoblast induction factor; F4/F80 – EGF-like module-containing mucin-like hormone receptor-like 1; Iba-1 – allograft inflammatory factor; CD68 – cluster of differentiation 68; TREM-2 – triggering receptor expressed on myeloid cells 2.

**Figure 4.**
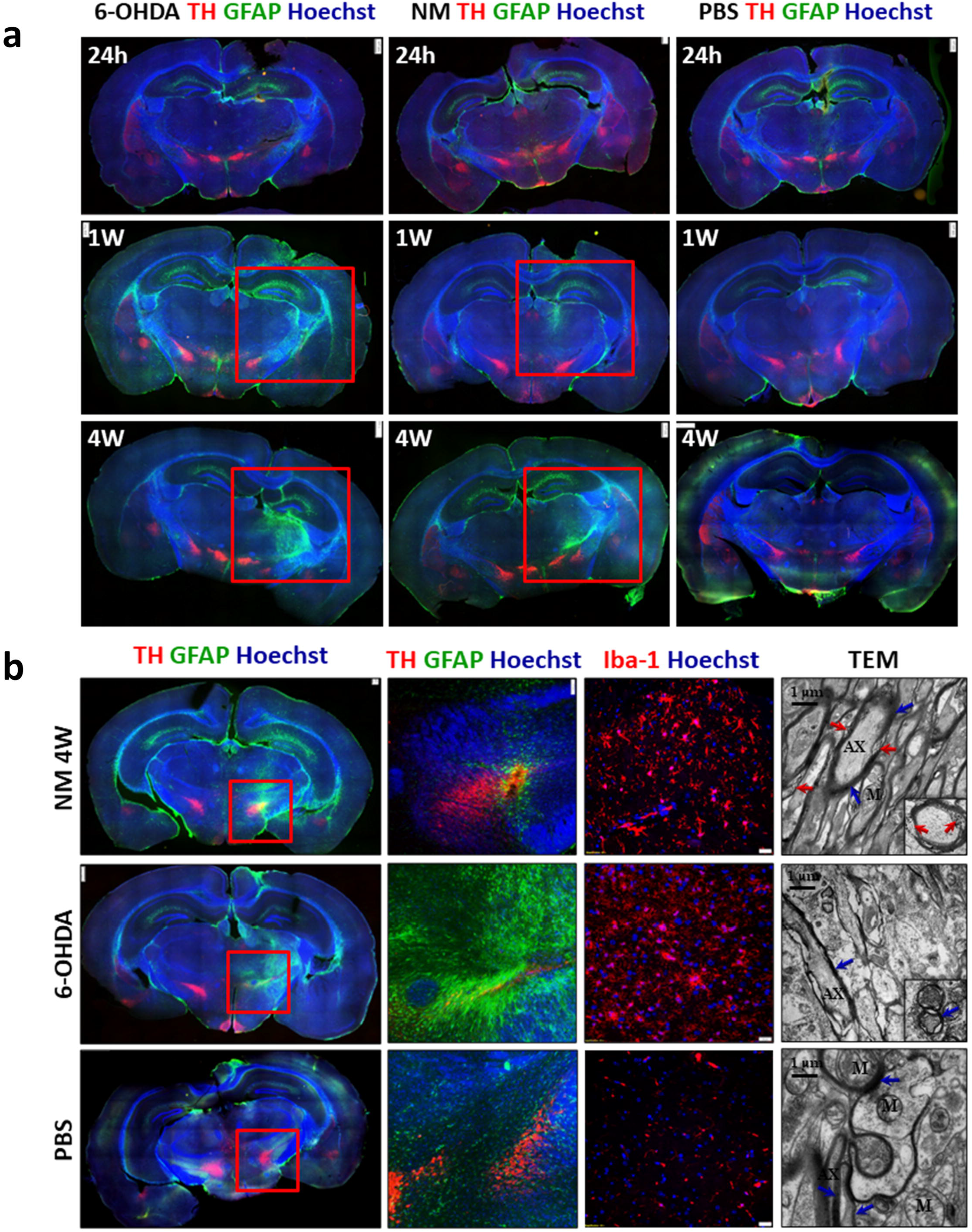
Synthetic neuromelanin activates microglia and astrocytes. Immunofluorescent staining and TEM micrographs of the brain (SNpc) after administration of 6-OHDA, NM and PBS. Cross-section of the brain. **a), b)** Marked fragments shows stained astrocytes – GFAP (green) and degrading dopaminergic neurons – TH (red). Iba-1 staining shows activation of microglia. Degradation of dopaminergic neurons and activation of microglia is not observed after PBS administration. Ultrastructure of mouse brain after administration of neuromelanin; 6-OHDA and PBS (control group) (**b**). After neuromelanin administration symptoms of demyelination are visible (red arrows). In 6-OHDA and PBS treated mice myelin sheaths have not shown signs of disintegration. AX - axon; M - mitochondria; blue arrows - myelin sheath; red arrows - sites of demyelination; TEM, bar=1µm.

### Neuromelanin and 6-OHDA injection activates in-vivo microglia and astrocytes contributing to the development of inflammation and damage to dopaminergic neurons in the substantia nigra

In order to visualize the effect of neuromelanin and 6-OHDA administered to the left hemisphere of the mouse brain on the protein level the immunofluorescence staining on the transverse sections of the brains were performed. For this purpose, 50 µm thick sections of brains obtained after fixation of the brains in PFA and those cut on a microtome were stained for the presence of GFAP, a marker of astrocytes, TH, an enzyme which is a marker of dopaminergic neurons and DAPI in order to visualize the cell nuclei. Additionally, in order to visualize Iba-1, a marker characteristic for microglia, 3 µm thick sections of brains fixed in 40% formalin and embedded in paraffin blocks were prepared. Figure 4 shows the immunofluorescence staining at the time points: 24h, 1 week (1W), 4 weeks (4W) for GAPDH, TH and DAPI, while Fig. 4b shows additionally the staining on Iba-1 4 weeks after injection and an electron microscope images 4 weeks after injection. 24 hours after injection of 6-OHDA, neuromelanin, and PBS do not induce GFAP expression or decrease in TH expression, while already 1 week after injection, a significant increase in GFAP expression is observed both after 6-OHDA injection and NM in the injection area in the hippocampus and in the substantia nigra region, this effect is not observed after PBS administration (Fig. 4a). 4 weeks after 6-OHDA injection there was a very strong expression of GFAP correlating with a decrease in TH amount, similar to NM, that effect was not observed in the case of PBS administration (Fig. 4a). Figure 4b shows brain sections stained with GFAP and TH 4 weeks after neuromelanin administration along with a zoom on substantia nigra showing damage to dopaminergic neurons manifested by a decrease in TH expression. This figure also shows the microglia visualized by increasing expression of the marker Iba-1. After 6-OHDA administration, astrocytosis was visible, correlating with a drastic decrease in TH level and severe microgliosis. After PBS administration, no increase in GFAP expression or decrease in TH expression and no increase in Iba-1 expression was observed (Fig. 4b). Changes in the morphology of the mitochondria after NM administration were observed and 6-OHDA compared to the PBS control in the photomicrographs taken with using the transmission electron microscope (TEM) (Fig. 4b). Mitochondria after NM administration became large compared to the mitochondria after PBS administration. Demyelination of neurons and accumulation of NM grains within the myelin after administration of neuromelanin was observed (Fig. 4b). Administration of 6-OHDA resulted in complete cell lysis, which makes ultrastructure of individual cell organelles indistinguishable. The above results indicate that the administration of NM disturbs cell function and causes chronic inflammation leading to decrease in the number of dopaminergic neurons, however, these changes are not as rapid and radical as in the case of administration of a strong neurotoxin 6-OHDA. Thanks to this, the newly obtained model reflects well certain aspects of PD and can be used to study changes in SNpc that accompany the death of dopaminergic neurons.

### Neuromelanin and 6-OHDA administration cause loss of dopaminergic neurons in the brain

Gradual loss of dopaminergic neurons at the RNA and protein level after administration of 6-OHDA as well as NM were confirmed by a behavioral spontaneous evaluation test behavior of the mouse in the cylinder (Fig. 5a). Spontaneous forelimb use was measured for 3 minutes to asses spontaneous mice behavior in the cylinder. U Mann Whitney test revealed significant effects in the amount of forelimb uses (forelimb steps) in both groups – with 6-OHDA and neuromelanin administration – from second week through whole observation in comparison with analogous timepoints in PBS group. U Mann Whitney test showed a significant reduction in the number of forelimb steps made by 6-OHDA mice compared with PBS mice in second week (p<0,0001), third week (p=0,0002) and fourth week (p<0,0001). The significant reduction in the number of forelimb steps was also revealed by U Mann Whitney test for the neuromelanin group in second, third and fourth week (p<0,0001, p=0,0075 and p=0,0002, respectively). In third and fourth week, the decrease in forelimb steps was higher for 6-OHDA group compared with neuromelanin-receiving group. These changes could be attributed to a radical damage of striatal dopaminergic neurons (>95% loss of striatal dopamine) caused by 6-OHDA administration (28). In order to measure long-term loss of dopaminergic neuron function, spontaneous activity in cylinder test was asses 1 and 27 weeks after neuromelanin or PBS administration. U Mann Whitney test revealed a significant decrease (p=0,0043) in the percentage of forelimb steps made by neuromelanin mice (Fig. 5b). There was not such tendency in PBS - receiving group.

**Figure 5.**
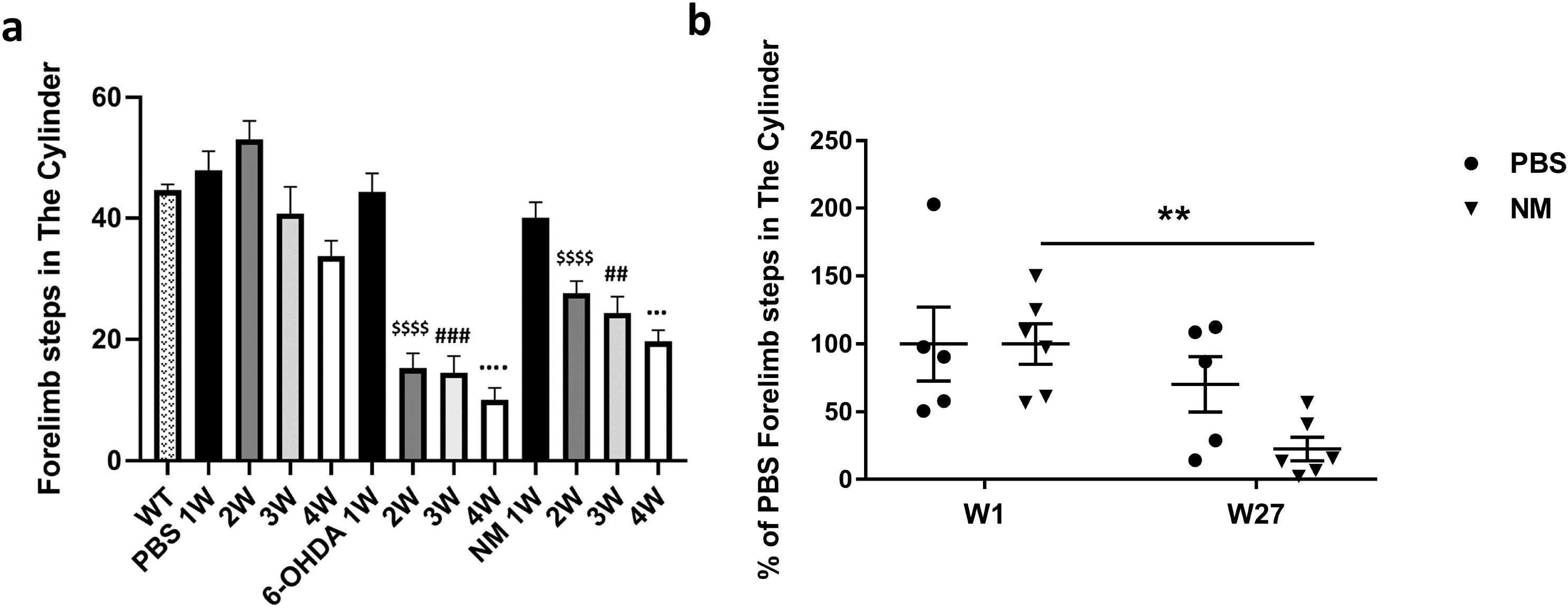
Administration of synthetic neuromelanin causes loss of dopaminergic neurons in the brain. **a)** Spontaneous activity in wild-type, PBS, 6-OHDA and neuromelanin (NM) treated mice in 1, 2, 3 and 4 weeks after administration. Values are expressed as mean ± SEM. Statistical analysis were performed used U Mann Whitney test. $ - compared to PBS week 2 (2W), # - compared to PBS week 3 (3W), – compared to PBS week 4 (4W). Two signs p<0,01, three signs p<0,001, four signs p<0,0001. **b)** Spontaneous activity in cylinder test in mice treated with PBS or neuromelanin (NM) 1 and 27 weeks after the administration. Values are expressed as mean ± SEM. Statistical analysis were performed used U Mann Whitney test. ** - compared to NM at week 1 (W1).

## Discussion

The aim of the research undertaken by our team was, on the one hand, to obtain a more physiological, mouse model of Parkinson’s disease induced by synthetic neuromelanin than previously used (7). On the other hand, we wanted to observe the response of immune cells such as microglia and astrocytes in the brain to the presence of synthetic neuromelanin. Leakage of neuromelanin from dopaminergic neurons in the substantia nigra has been proposed as a possible mechanism in the development of Parkinson’s disease (34). It is thought that the leakage of neuromelanin could trigger an immune response in the brain, leading to inflammation and oxidative stress (7, 34). This inflammation and oxidative stress could then contribute to the degeneration of dopaminergic neurons and the subsequent loss of dopamine in the brain, which is a hallmark of Parkinson’s disease (14, 35, 36). In addition, the accumulation of neuromelanin outside of the neurons could also cause toxicity by generating reactive oxygen species (ROS) (35).

The exact mechanisms underlying the leakage of neuromelanin from neurons are not yet fully understood, but several theories have been proposed (10). Age-related degradation, oxidative stress, inflammation and genetic factors may affect the protective mechanisms that prevent the leakage of neuromelanin, making individuals more susceptible to the development of Parkinson’s disease (10, 12).

In this study we have shown that the administration of synthetic neuromelanin, which activates astrocytes and microglia, induces the inflammation and may be involved in degeneration of dopaminergic neurons. Although the central nervous system (CNS) is considered an immune-privileged tissue, it can initiate an immune response through microglia, resident CNS tissue macrophages that can be activated by various stimuli against pathogens or endogenous threat signals (37, 38, 39). If the mechanisms involved in terminating inflammation are insufficient, chronic inflammation may develop It may also occur in response to molecules secreted by degenerating neurons, a condition called neuroinflammation, which plays a key role in neurodegenerative diseases (34, 39). In the pathogenesis of PD, the integrity of the blood-brain barrier (BBB) is compromised and components of the innate immune system may be activated allowing the recruitment and activation of the adaptive immune system (40). This allows pro-inflammatory molecules to reach the central nervous system (CNS) from the circuit passing through the BBB. The most numerous astrocytes in the CNS are functionally connected to the BBB, receiving signals from the periphery and the interior of the CNS (39). Astrocytes participate in the maintenance and permeability of the BBB and are key regulators of neuronal activity and cerebral blood flow (41). In vitro and in vivo studies demonstrate the important role that astrocytes play in the neuroinflammatory processes in PD. Astrocytes, like microglia, respond to inflammatory stimuli such as IL-1β, LPS and TNF-α by producing more pro-inflammatory cytokines (14). Reactive astrogliosis has been reported in the affected brain areas of PD (36) patients, indicating the possible involvement of astrocytes in the immune response in PD. It has been shown that LPS-responsive astrocytes can be detrimental due to the upregulation of genes for the classical complement cascade, which is believed to cause loss of synapses and neuronal loss in neurodegenerative diseases. Reactive astrocytes induced by neuritis are referred to as A1 reactive astrocytes, and those induced by ischemia are referred to as A2 reactive astrocytes (23,41). We have shown that synthetic neuromelanin induces strong astrocytosis. The pro-inflammatory mediators released by activated astrocytes act on their cognate receptors expressed in microglia and further increase microglial activation, bringing them to an over-activated state. Microglia are immune cells of the CNS, constituting 5-10% of all brain cells (17, 42). Microglia are at rest in the absence of any stimulus, which is achieved by the immunosuppressive microenvironment present in the CNS, where immunoregulatory molecules are expressed and/or released by healthy neurons. Resting microglia can be activated in two classic phenotypes, M1 and M2 (43), depending on effector signals from its microenvironment. Microglia upregulate markers of inflammation on the cell surface, including MHC class I and II, and cytokine and chemokine receptors. In the presence of LPS and IFN-γ, microglial cells polarize to the M1 phenotype and secrete pro-inflammatory cytokines that contribute to the dysfunction of dopaminergic neurons (neurodegeneration). We have shown that the activation of microglia, synthetic neuromelanin, is correlated with a decrease in TH and a loss of dopaminergic neurons. Short-term activation of microglia is thought to be neuroprotective, while chronic activation is considered a potential mechanism in neurodegenerative diseases (23).

In conclusion, synthetic neuromelanin has been used in animal models as a tool to study the role of neuromelanin in Parkinson’s disease and to evaluate potential treatments for the disease. These animal models have contributed to our understanding of the underlying mechanisms of Parkinson’s disease and have helped to advance the development of new and more effective treatments for the disease.

## Acknowledgments

We would like to thank O. Woźnicka form the Department of Cell Biology and Imaging, Institute of Zoology and Biomedical Research, Jagiellonian University, for her skilled technical assistance with TEM material preparation and observations.

